# Loss of *OsARF18* confers glufosinate ammonium herbicide resistance in rice

**DOI:** 10.1101/2023.06.05.543806

**Authors:** Jin-Qiu Xia, Da-Yu He, Qin-Yu Liang, Zheng-Yi Zhang, Jie Wu, Zi-Sheng Zhang, Jing Zhang, Ping-Xia Zhao, Cheng-Bin Xiang

**Affiliations:** Division of Life Sciences and Medicine, Division of Molecular & Cell Biophysics, Hefei National Science Center for Physical Sciences at the Microscale, University of Science and Technology of China, The Innovation Academy of Seed Design, Chinese Academy of Sciences, Hefei, Anhui Province 230027, China

**Keywords:** rice (*Oryza sativa* L.), glufosinate ammonium, herbicide resistance, *gar1*, *OsARF18*, salt and osmotic stress

## Abstract

Weed is one of the major biotic stresses that causes severe loss of crop yield. Herbicide is one of the most cost-effective ways to control weeds. Thus, the development of herbicide-resistant crops is critical for the application of herbicides. To isolate new glufosinate ammonium resistance loci, we screened a rice ethyl methyl sulfonate-mutagenized library and obtained the *g*lufosinate *a*mmonium-*r*esistant mutant *gar1-1*. *GAR1* encodes auxin response factor 18 (*OsARF18*). A G-to-A substitution in the coding region of *OsARF18* results in loss of function of *OsARF18* and thereby enhances glufosinate ammonium resistance of *gar1-1*, which was confirmed by three additional CRISPR/Cas9-edited *gar1* alleles. *GLUTAMINE SYNTHETASE 1;1* (*OsGS1;1*) and *GLUTAMINE SYNTHETASE 1;2* (*OsGS1;2*) were upregulated in *gar1-1* upon glufosinate ammonium treatment, directly contributing to increased GS activity that enhances glufosinate ammonium herbicide resistance. We further show that OsARF18 suppresses *OsGS1;1* and *OsGS1;2* expression.

Comparative transcriptomic analyses reveal a huge shift in the gene expression profile involved in stress tolerance and growth. A large number of detoxification-related genes are enriched in *gar1-1*, which may also contribute to enhanced herbicide resistance. Moreover, stress tolerance-related genes are upregulated and growth-related genes are downregulated in *gar1-1*, consistent with the improved tolerance to salt and osmotic stress of *gar1* mutants. Taken together, our study demonstrates that *OsARF18* is a negative regulator of glufosinate ammonium resistance as well as salt and osmotic stress tolerance, suggesting a role in balancing the stress response and growth.

## INTRODUCTION

Weed is one of the most common biotic stresses causing severe crop yield decline. Weeds compete with crops for resources such as sunlight, water, fertilizer and growth space and serve as hosts of pests and diseases, which further reduces crop yield (Jin et al., 2022). Chemical herbicides are an economical and effective method for weed control and are widely used in modern agriculture (Amna et al., 2019). Over the past decades, the use of chemical herbicides has increased dramatically with the introduction of large numbers of genetically modified crops, including corn, soybeans, and cotton. Therefore, it is of great significance to develop crops resistant to new herbicides to alleviate weed infestations and maintain sustainable crop production.

Glufosinate, as a broad spectrum, nonselective herbicide, has been widely used in the past few decades to control annual and perennial weeds (Takano and Dayan, 2020). Glufosinate is a structural analog of glutamic acid and specifically inhibits the activity of glutamine synthetase (GS), which is vital for catalyzing the synthesis of glutamine from glutamic acid and ammonium. Glutamate and glutamine are the initial products of plant nitrogen (N) assimilation from inorganic N to organic N, thus disrupting the process of N assimilation, causing excessive accumulation of ammonium and reactive oxygen species (ROS) and subsequent lipid peroxidation in organisms, inhibiting photosynthetic activity, and leading to cell death (Droge-Laser. et al., 1994; Takano and Dayan, 2020). The evolution rate of glufosinate-resistant weeds is relatively slow, and it is rare for weeds to gain cross-resistance with other herbicides. To date, only *Amaranthus palmeri*, *Eleusine indica*, *Poa annua*, and *Lolium perenne* have been reported to have evolved resistance to glufosinate (Heap, 2022). Therefore, the development of crops resistant to glufosinate herbicide is of great significance for weed control, especially for direct seeding and rain-fed areas.

The glufosinate herbicide resistance genes *BAR* and *PAT* were cloned from *Streptomyces hygroscopicus* and *Streptomyces viridochromogenes Tü494*, respectively, and identified to confer glufosinate resistance by detoxifying the herbicide (Strauch et al., 1988; Thompson et al., 1987), and these genes have been widely used to develop glufosinate-resistant crops. Modification of endogenous targets of glufosinate in plants is also considered to be a useful approach to improve glufosinate resistance. Yan et al. effectively induced A-G nucleotide conversion in the *OsGS1* gene by using new TadA variant technology, thus obtaining glufosinate-resistant rice (Yan et al., 2021). Target site resistance of glufosinate in plants is generally thought to be caused by specific mutations in *GS* genes (Avila Garcia et al., 2012; Donald James. et al., 2018; Noguera et al., 2022; Tian et al., 2015). The mutated GS no longer binds glufosinate and hence confers herbicide resistance. Moreover, co-overexpression of *OsGS1;1* and *OsGS2* also confers enhanced glufosinate resistance (James et al., 2018). It is unclear whether other glufosinate resistance mechanisms exist in nature or whether new resistance loci can be generated through mutagenesis.

Auxin response factors (ARFs) are a class of plant-specific transcription factors that play a vital role in plant growth, development and response to biotic and abiotic stresses by binding to TGTCGG/TGTCNN *cis*-element in the promoter region of downstream target genes (Guilfoyle, 2015; Guilfoyle and Hagen, 2007; Li et al., 2022b; Song et al., 2022; Truskina et al., 2020). For instance, *OsARF4/6/11/18/25* play key roles in regulating grain and yield by affecting grain size and shape (Hu et al., 2018; Huang et al., 2016; Qiao et al., 2021; Sims et al., 2021; Zhang et al., 2018). *OsARF11* also strongly influences plant height, tiller numbers, root system and flag leaf angle (Liu et al., 2018; Sims et al., 2021). In the context of biotic and abiotic resistance, *OsARF12* and *OsARF16* positively regulate rice-dwarf virus defense, whereas *OsARF11* and *OsARF5* have the opposite effect (Qin et al., 2020). Overexpression of *ZmARF4* enhanced root morphogenesis under low phosphorus stress and improved salinity and osmotic stress tolerance (Li et al., 2022a). *SlARF10* participates in water transport by regulating stomatal development and aquaporin expression (Liu et al., 2016). *miR160-ARF18* mediates salt stress tolerance in peanut (Tang et al., 2022). OsARF23 plays an important role in the plant response to drought stress, which possibly alters the root growth angle by inhibiting *DEEPER ROOTING 1* expression, thus reducing plant drought resistance (Uga et al., 2013). *OsARF18* has been reported to be involved in multiple physiological processes, including negatively regulating salt tolerance and nitrogen use efficiency in rice by modulating aspartic acid production and NH_4_^+^ accumulation, and the OsARF18-OsARF2-OsSUT1 module mediates the auxin signaling cascade to regulate sourceLsink carbohydrate partitioning and reproductive organ development (Deng et al., 2022; Huang et al., 2016; Zhao et al., 2022). Furthermore, the transcriptional activity of OsARF18 is regulated by the acetylation of histone H3 in callus by OsHDA710, reducing callus formation of the rice mature embryo by repressing *PLT1* and *PLT2* (Zhang et al., 2020). However, there have been no reports of *ARF* genes involved in herbicide resistance in plants.

In this study, we screened an ethyl methyl sulfonate (EMS)-mutagenized rice library with glufosinate herbicide and isolated a glufosinate-resistant mutant named *glufosinate ammonium resistance 1-1* (*gar1-1*). *GAR1* encodes the transcription factor OsARF18. Knockout of *OsARF18* confers enhanced glufosinate resistance as well as improved tolerance to salt and osmotic stress. Our results reveal that loss of *OsARF18* derepresses the expression of *OsGS1;1* and *OsGS1;2*, enhancing GS activity in *gar1* mutants and thus promoting glufosinate herbicide resistance, and upregulates the expression of a large number of stress tolerance– and detoxification-related genes that may contribute to enhanced glufosinate resistance and improved tolerance to salt and osmotic stress. Therefore, our study reveals a novel genetic locus of glufosinate herbicide resistance and provides a potential candidate gene for the development of glufosinate– and abiotic stress-tolerant crops.

## RESULTS

### Isolation and characterization of *gar1-1* with enhanced glufosinate ammonium resistance

To isolate glufosinate ammonium resistance mutants, we carried out a genetic screen of the EMS-mutagenized M_2_ seeds in the Longgeng31 wild-type (LG31) background. The mutagenized library consists of 103 pools, and each pool is composed of the bulk M_2_ seeds collected from approximately 1000 individual M_1_ plants. Two-week-old M_2_ seedlings were sprayed with 2 g/L glufosinate herbicide (Ruikai Chemical, Shijiazhuang, China). After 7-10 days, very few surviving seedlings were collected as putative mutants and named *glufosinate ammonium resistance* (*gar*). After rescreening the progeny of *gar* mutants with 2 g/L glufosinate herbicide, we confirmed the glufosinate resistance phenotype of several *gar* mutants, including *gar1-1*.

Figure 1A shows that two-week-old *gar1-1* seedlings exhibited significantly enhanced resistance after being sprayed with 2 g/L glufosinate, whereas all wild-type seedlings died (Figure 1A). To evaluate the *gar1-1* glufosinate resistance level, two-week-old seedlings of both wild-type LG31 and *gar1-1* were sprayed with glufosinate at different concentrations (0, 2, 4, 6, 8 and 10 g/L). All *gar1-1* seedlings survived 10 g/L glufosinate, the highest concentration challenged, despite part of the leaves turning yellow and growth being significantly inhibited, whereas the wild-type LG31 did not survive 2 g/L glufosinate, the lowest concentration challenged (Figure 1B). The results clearly show that *gar1-1* is significantly more tolerant to glufosinate than wild-type LG31.

**Figure 1.**
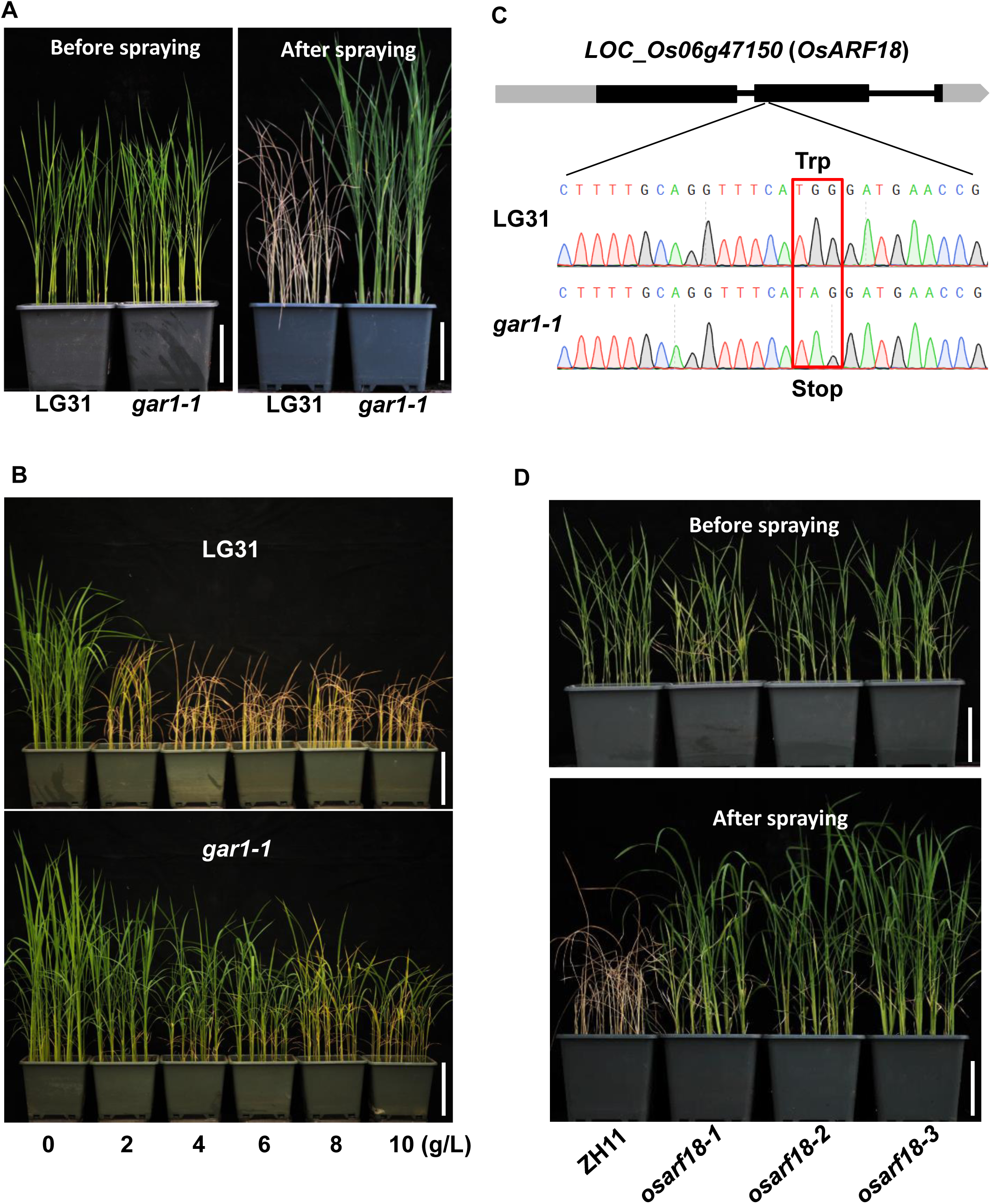
Glufosinate ammonium herbicide resistance phenotype of *gar1-1*. **A.** Glufosinate resistance phenotype of the *gar1-1* mutant. Seedlings of LG31 and *gar1-1* grown in the soil for 2 weeks before (left) and after being sprayed with 2 g/L glufosinate (right). The image was recorded before and 10 days after the seedlings were sprayed with glufosinate. **B.** Evaluation of *gar1-1* resistance to glufosinate. Two-week-old LG31 and *gar1-1* seedlings were sprayed with glufosinate at different concentrations (0, 2, 6, 8, and 10 g/L). The images were recorded 10 days after herbicide application. **C.** Schematic representation of *LOC_Os06g47150* (*OsARF18*) and identification of the *gar1-1* mutation by sequencing. Gray and black boxes represent untranslated regions (UTRs) and exons, respectively, and black lines denote introns. The red box shows the codon change from TGG to TAG, resulting in a nonsense mutation. Bar= 10 cm. **D.** Glufosinate resistance phenotype of *OsARF18* knockout mutants created by the CRISPR/Cas9 technique. Seedlings of ZH11, *osarf18-1*, *osarf18-2* and *osarf18-3* lines grown in the soil for 2 weeks before (upper) and after spraying with 2 g/L glufosinate (lower). The images were recorded before and 10 days after the seedlings were sprayed with the herbicide.

### Identification of the mutation in *gar1-1*

To identify the mutation in *gar1-1*, we backcrossed the mutant with LG31 wild type to generate the BC1F1 population. BC1F1 progeny were selfed to generate the BC1F2 population. Both the BC1F1 and BC1F2 populations were phenotyped for glufosinate resistance. All BC1F1 progeny were sensitive to glufosinate, whereas the BC1F2 progeny showed a 229/66 ratio of glufosinate sensitive/resistant phenotype [χ^2^ =1.08<χ^2^(0.05) =3.84], near a 3:1 segregation ratio, indicating that *gar1-1* is caused by a single recessive nuclear gene mutation. Then, we bulked DNA from 20 BC1F2 progeny with a glufosinate-resistant phenotype for whole-genome sequencing. Based on the MutMap method, we identified a point mutation in the *LOC_Os06g47150* locus, which encodes the auxin response factor OsARF18. A G-to-A substitution at the 1133^th^ nucleotide in the coding region of *OsARF18* caused a nonsense mutation from tryptophan (TGG) to a stop codon (TAG), which resulted in the loss of function of *OsARF18* (Figure 1C).

### Confirmation of loss of *OsARF18* causing glufosinate resistance

To confirm whether loss of *OsARF18* causes a glufosinate resistance phenotype, we generated three *OsARF18* knockout mutants (*osarf18-1, osarf18-2* and *osarf18-3*) using the CRISPR/Cas9 genome editing system in the wild-type Zhonghua11 (ZH11) background (Figure S1A-C) (Lu et al., 2017). Then, all the knockout lines (*osarf18-1, osarf18-2* and *osarf18-3*) and wild-type ZH11 were assayed for glufosinate resistance under soil growth conditions. Under normal growth conditions, there were no obvious differences between ZH11 and the knockout mutants. However, all the knockout mutants showed a glufosinate resistance phenotype similar to *gar1-1* 10 days after being sprayed with 2 g/L glufosinate herbicide, displaying a significantly enhanced glufosinate resistance phenotype with 100% survival, whereas all the wild-type ZH11 seedlings died (Figure 1D).

Taken together, our results reveal that *OsARF18* is a negative regulator of rice glufosinate resistance, a genetic locus of glufosinate herbicide resistance that has not been previously reported.

### The expression pattern of *OsARF18/GAR1* in response to abiotic stresses

To determine the expression pattern of *OsARF18*, we generated *OsARF18pro:GUS* transgenic plants. GUS staining results showed that *OsARF18* was expressed in all tissues detected (Figure 2A-J). In addition, *OsARF18* transcript levels were measured by RTLqPCR in the indicated tissues of wild-type ZH11 plants, with higher levels in seedling shoots (Figure 2K). To further investigate the response of *OsARF18* expression to abiotic stresses, two-week-old ZH11 seedlings were treated with NaCl, PEG4000, H_2_O_2_, ABA, CdCl_2_, and 4°C conditions. The RTLqPCR analysis results showed that *OsARF18* expression was slightly induced by abiotic stresses (Figure 2L), suggesting that *OsARF18* may be involved in stress responses.

**Figure 2.**
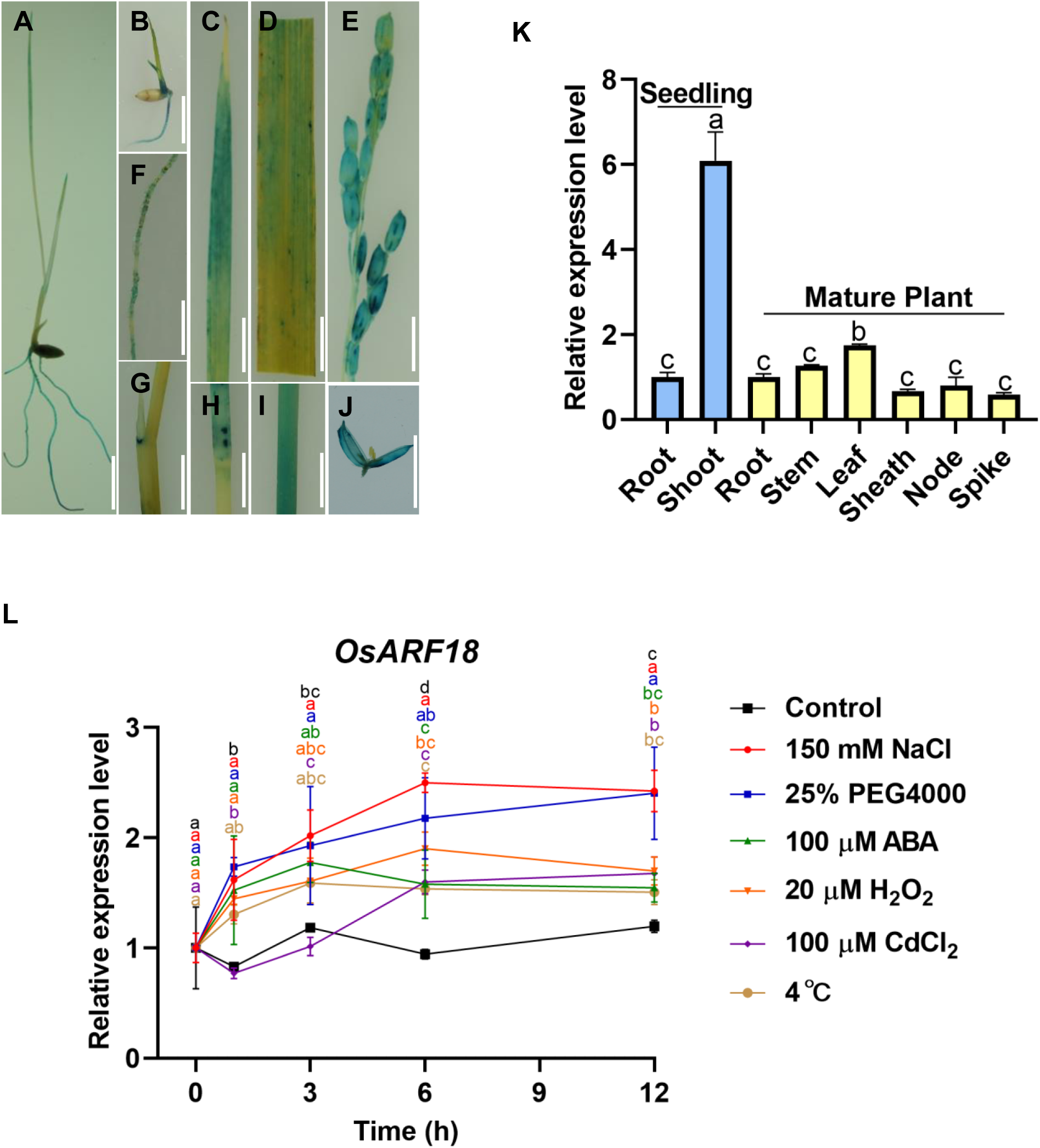
Expression pattern of *OsARF18* and response to abiotic stresses. **A-J**. GUS staining of various tissues of *OsARF18pro:GUS* transgenic plants. Seven– day-old seedlings (A), seeds germinating for 3 days (B), young leaves (C), mature leaves (D), panicles (E), roots (F), sheaths (G), nodes (H), stems (I) and caryopses (J). Bar = 1 cm. **K**. Expression of *OsARF18* in different tissues of two-week-old seedlings (roots and shoots) and at the flowering stage (root, stem, leaf, sheath, node and spike) of ZH11 wild type. *OsACTIN1* was used as an internal control. Values are the mean ± SD (n = 3). Different letters indicate significant differences by one-way ANOVA (p<0.05). **L.** Two-week-old seedlings of wild-type ZH11 grown in hydroponic culture were treated with 150 mM NaCl, 25% PEG4000, 100 μM ABA, 20 μM H_2_O_2_, and 100 μMCdCl_2_ in the culture medium and 4 L for the indicated times (0, 1, 3, 6, and 12 h). RNA was extracted for RTLqPCR analyses. *OsACTIN1* was used as an internal control. Values are the mean ± SD (n = 3). Different letters indicate significant differences by one-way ANOVA (p<0.05).

### Subcellular localization of OsARF18

To analyze the subcellular localization of OsARF18, we generated a *35Spro:OsARF18-eGFP* fusion construct and transiently expressed the *35Spro:eGFP* and *35Spro:OsARF18-eGFP* fusion constructs in the leaves of *Nicotiana benthamiana*. The green fluorescence signals of the control vector (*35Spro:eGFP*) were distributed throughout the whole cell (Figure 3A), whereas the fluorescence signals of the OsARF18-eGFP fusion protein were detected only in the nucleus (Figure 3B). Moreover, we also generated stable *35Spro:OsARF18-eGFP* transgenic rice lines, which showed the fluorescent signals of OsARF18-eGFP in the nucleus of rice roots (Figure 3C).

**Figure 3.**
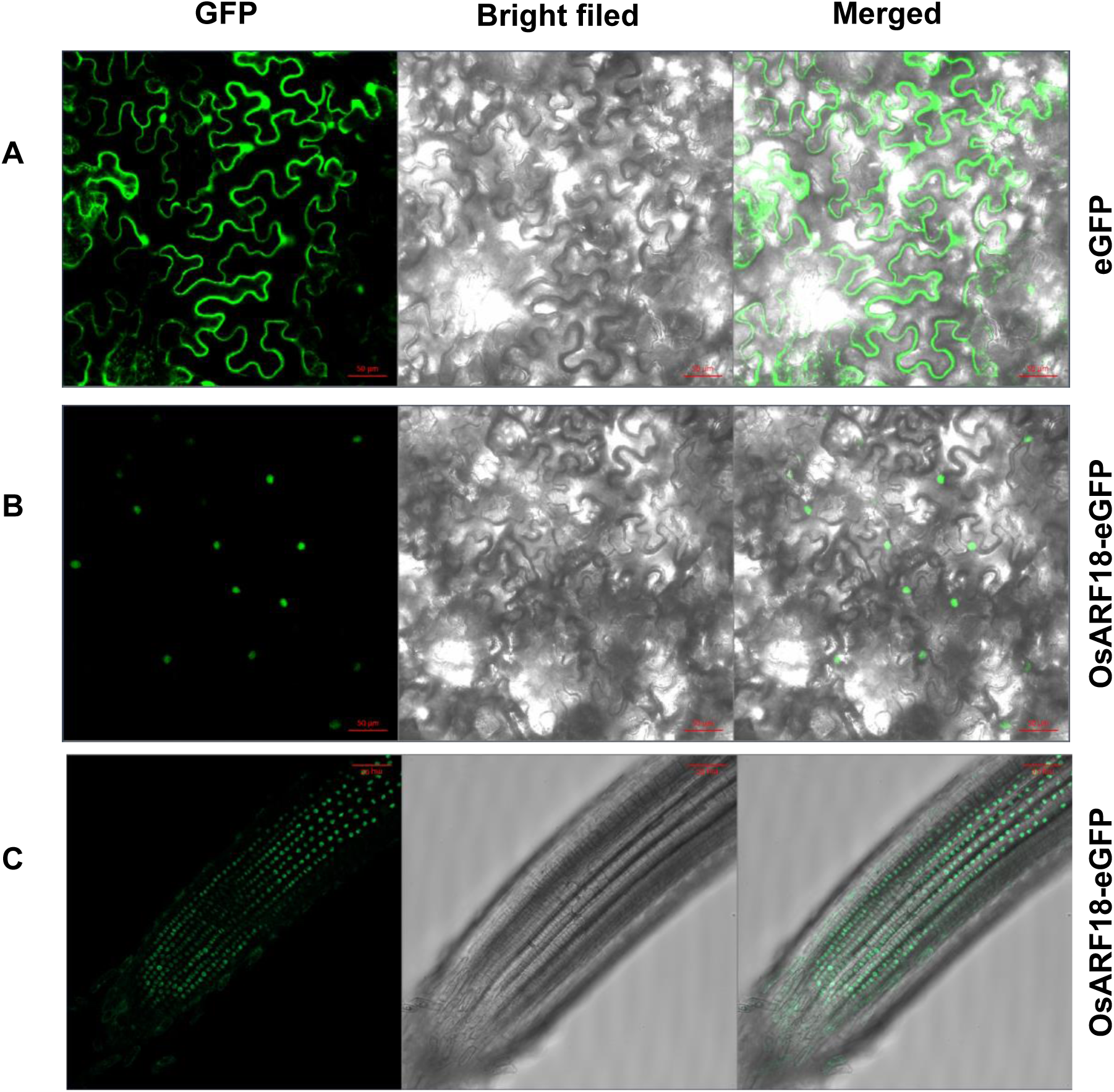
Subcellular localization of OsARF18. **A-B.** Fluorescence signals of *35Spro:eGFP* (eGFP) **(A)** and *35Spro:OsARF18-eGFP* (OsARF18-eGFP) **(B)** plasmids that were transiently expressed in epidermal cells of tobacco leaves. **C.** Fluorescence signals in the root tips of *35Spro:OsARF18-eGFP* transgenic lines. Bar = 50 μm.

### Enhanced GS activity contributing to enhanced herbicide resistance

To explore the mechanism by which *OsARF18* negatively regulates glufosinate resistance, we wondered whether GS capacity was enhanced in the mutant. Thus, we measured the transcript abundance of *OsGS1;1* and *OsGS1;2* and found that it was higher in the mutants than in the wild type when the seedlings were treated with glufosinate ammonium (Figure 4A and 4B). Likewise, GS activity was also increased significantly in the *gar* mutants (Figure 4C). These results indicate that increased GS activities due to upregulated expression of *OsGS1;1* and *OsGS1;2* contribute to enhanced glufosinate resistance, consistent with the phenotype of *gar1* mutants.

**Figure 4.**
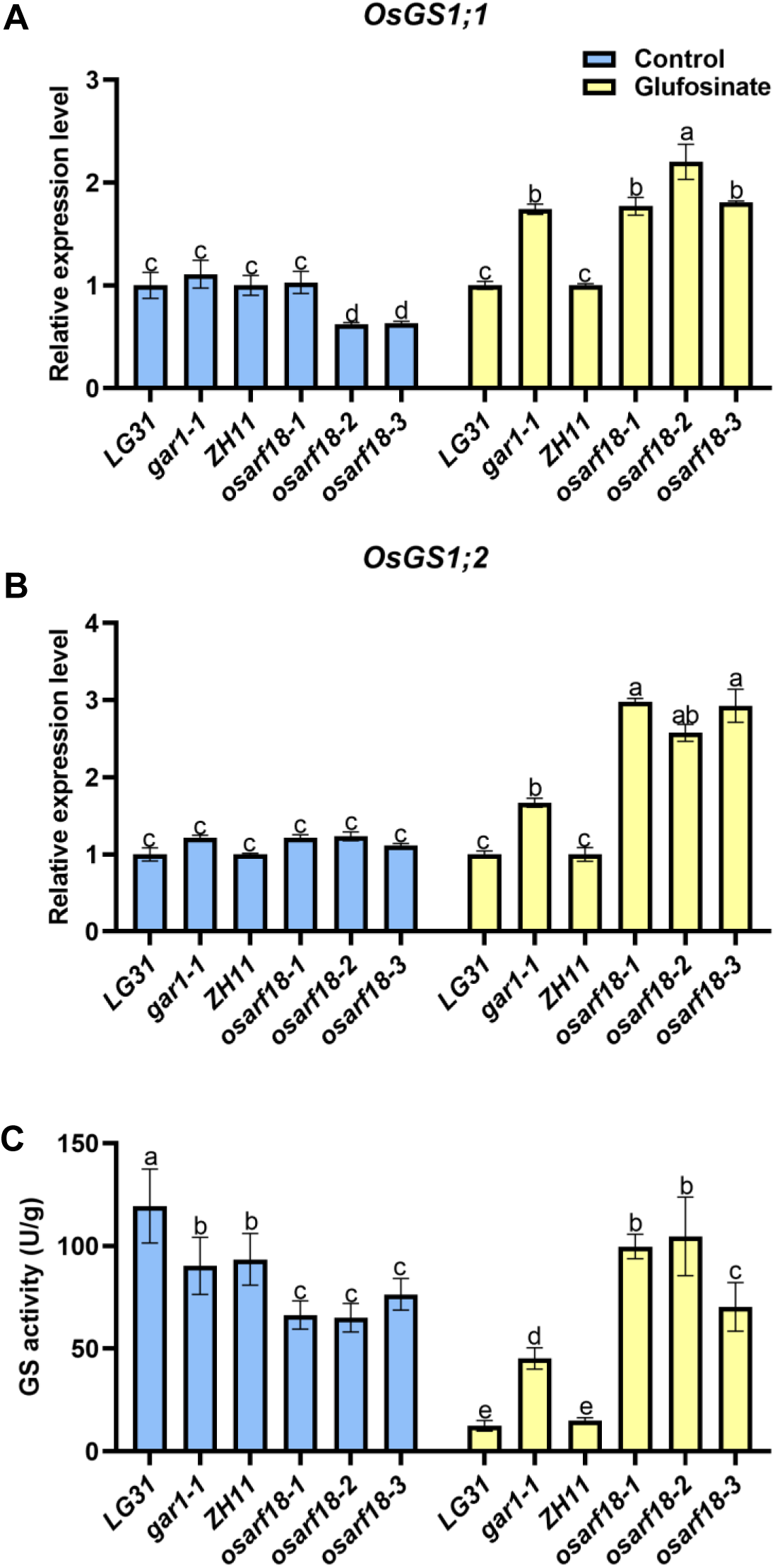
Transcript levels of *OsGS1;1* and *OsGS1;2* and GS enzyme activities were increased in *gar1* mutants. **A-B**. Expression levels of *OsGS1;1* (A) and *OsGS1;2* (B). Two-week-old seedlings of LG31, *gar1-1*, ZH11, *osarf18-1*, *osarf18-2,* and *osarf18-3* were sprayed with water (Control) or 2 g/L glufosinate (Glufosinate). Two days later, RNA was extracted from aerial parts for RTLqPCR analyses. *OsACTIN1* was used as an internal control. Values are the mean ± SD (n = 3). Different lowercase letters indicate significant differences (P < 0.05; one-way ANOVA). **C.** Glutamine synthetase activity. Two-week-old seedlings of LG31, *gar1-1*, ZH11, *osarf18-1*, *osarf18-2,* and *osarf18-3* were sprayed with water (Control) or 2 g/L glufosinate (Glufosinate). Two days later, GS activities were assayed as described in the methods. Values are the means ± SD (n = 3). Different lowercase letters indicate significant differences (P < 0.05; one-way ANOVA).

As a transcription repressor, OsARF18 may directly suppress the expression of *GS* genes. We searched the promoter sequences of *OsGS1;1* and *OsGS1;2* and found candidate ARF-binding cis elements (Figure 5A). ChIP-qPCR assays using *35Spro:OsARF18-eGFP* transgenic plants and anti-GFP antibodies showed significant enrichment of the *OsGS1;1* promoter region with the *cis1*-element and the *OsGS1;2* promoter region with the *cis1*-element and *cis2-*element compared with the control (Figure 5B). This result indicates that OsARF18 binds to the TGTCTC/TGTCNN *cis*-element in the *OsGS1;1* and *OsGS1;2* promoters to regulate their expression, and a transient trans repression assay in tobacco leaves also showed that cotransfection of the effector *35Spro:OsARF18* and reporter *OsGS1;1pro:LUC* or *OsGS1;2pro:LUC* in *N. benthamiana* leaves produced significantly less LUC activity than the control (Figure 5C). These results indicate that OsARF18 represses the expression of *OsGS1;1* and *OsGS1;2 in vivo*.

**Figure 5.**
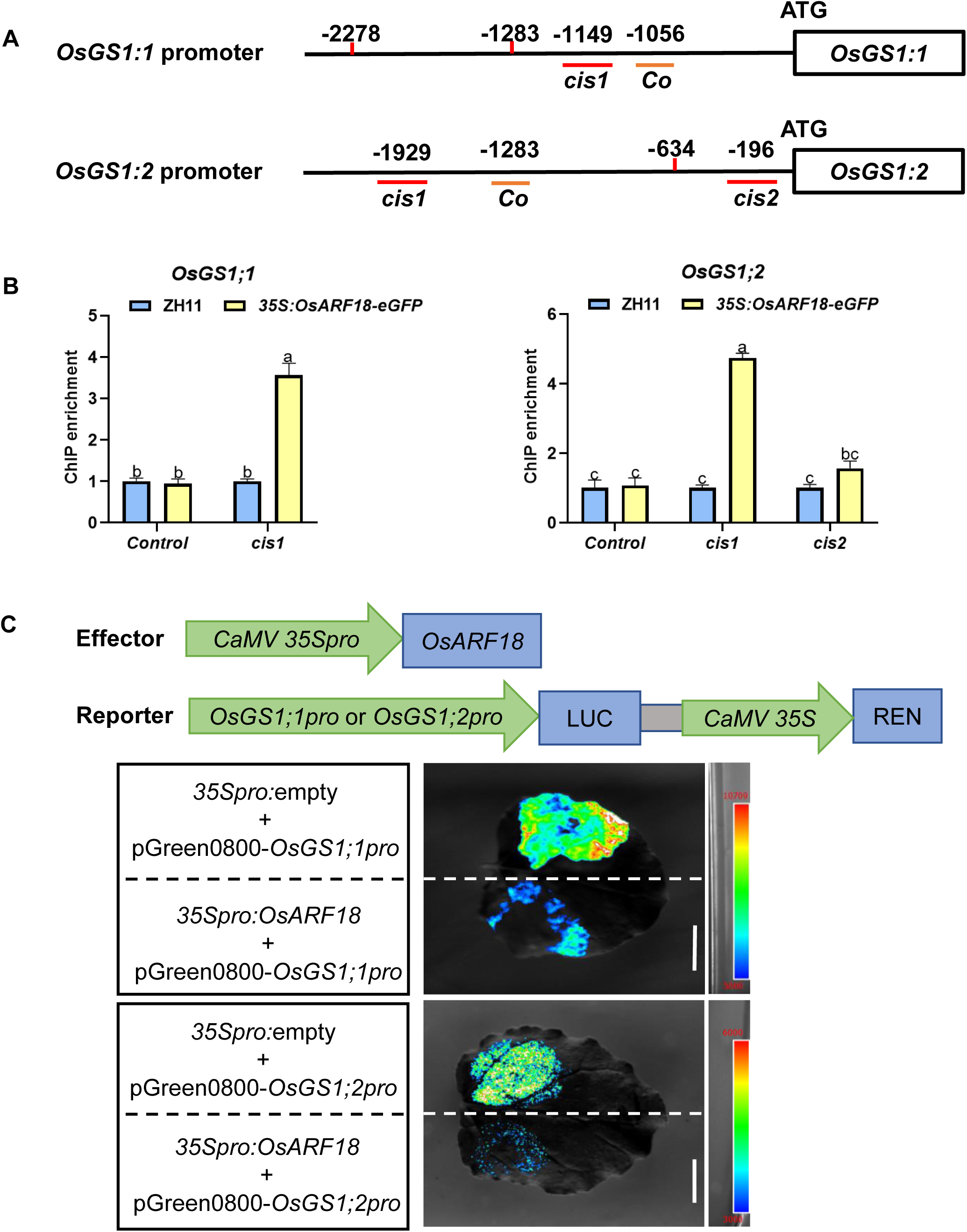
OsARF18 represses *OsGS1;1* and *OsGS1;2* expression by binding to the TGTCTC/TGTCNN motif in its promoter. **A.** Schematic illustration of the *OsGS1;1* and *OsGS1;2* promoter regions. Horizonal short color lines represent the sites of the predicted OsARF18-binding cis elements (red) or control sequence (orange). **B.** ChIP-qPCR assay. Two-week-old *35Spro:OsARF18-eGFP* plants and wild type plants (Control) were used for the ChIP-qPCR assay. Different letters denote significant differences evaluated by ANOVA with Tukey’s test (p<0.05). **C.** Transient trans repression assay. Repression of *OsGS1;1* and *OsGS1;2* expression by OsARF18 was determined by coinfiltrating tobacco leaves with *Agrobacterium* strains harboring the effector construct *35Spro:OsARF18* and *LUC* reporter driven by the *OsGS1;1* or *OsGS1;2* promoter. *35Spro:*empty, empty effector plasmid as control. pGreen0800, *LUC* reporter plasmid. Bar = 1 cm.

### The regulatory network of OsARF18 revealed by comparative transcriptomic analyses

To explore the regulatory network of OsARF18 as a transcriptional repressor, we performed high-throughput transcriptome deep sequencing analyses using shoots of two-week-old LG31 and *gar1-1* seedlings. We identified a total of 1704 differentially expressed genes (DEGs) (> 1.5-fold change, P < 0.05) in *gar1-1* compared with LG31, including 851 upregulated and 853 downregulated genes (Figure S2A).

To further analyze the transcription profiles of OsARF18-regulated DEGs, we used hierarchical clustering to analyze the DEGs between LG31 and *gar1-1*. As shown in Figure 6A, a number of genes associated with detoxification were enriched, including cytochrome P450s (*CYP707A5*, *CYP94C1*, *CYP76M1*, *CYP94C4*, *CYP96B4*, and *CYP71Z6*), glutathione S-transferases (*GST3, TCHQD1, GSTU16*), ABC transporters (*ABCG14*, *ABCG40*, *PDR4*, *PDR5*, and *PDR18*), and MATE transporter (*DTX49*). The transcript levels of genes encoding peroxidases (*PRX17*, *PRX19*), NADP-dependent oxidoreductase (*Os04g0497000*), and glutaredoxin (*GRX6*) in antioxidation were also increased in *gar1-1*. Moreover, several glutamate/glutamine-responsive genes, including *ERF109*, *WRKY53*, *bHLH13*, and *bHLH35*, were upregulated in *gar1-1*, which were possibly involved in nitrogen metabolism and stress responses and reflected an increased GS capacity and glutamate/glutamine levels, consistent with previous reports (Kan et al., 2015; Kan et al., 2017). Together, these results suggest that increased GS activity, detoxification, and antioxidation may collectively contribute to enhanced glufosinate resistance in *gar1-1*.

**Figure 6.**
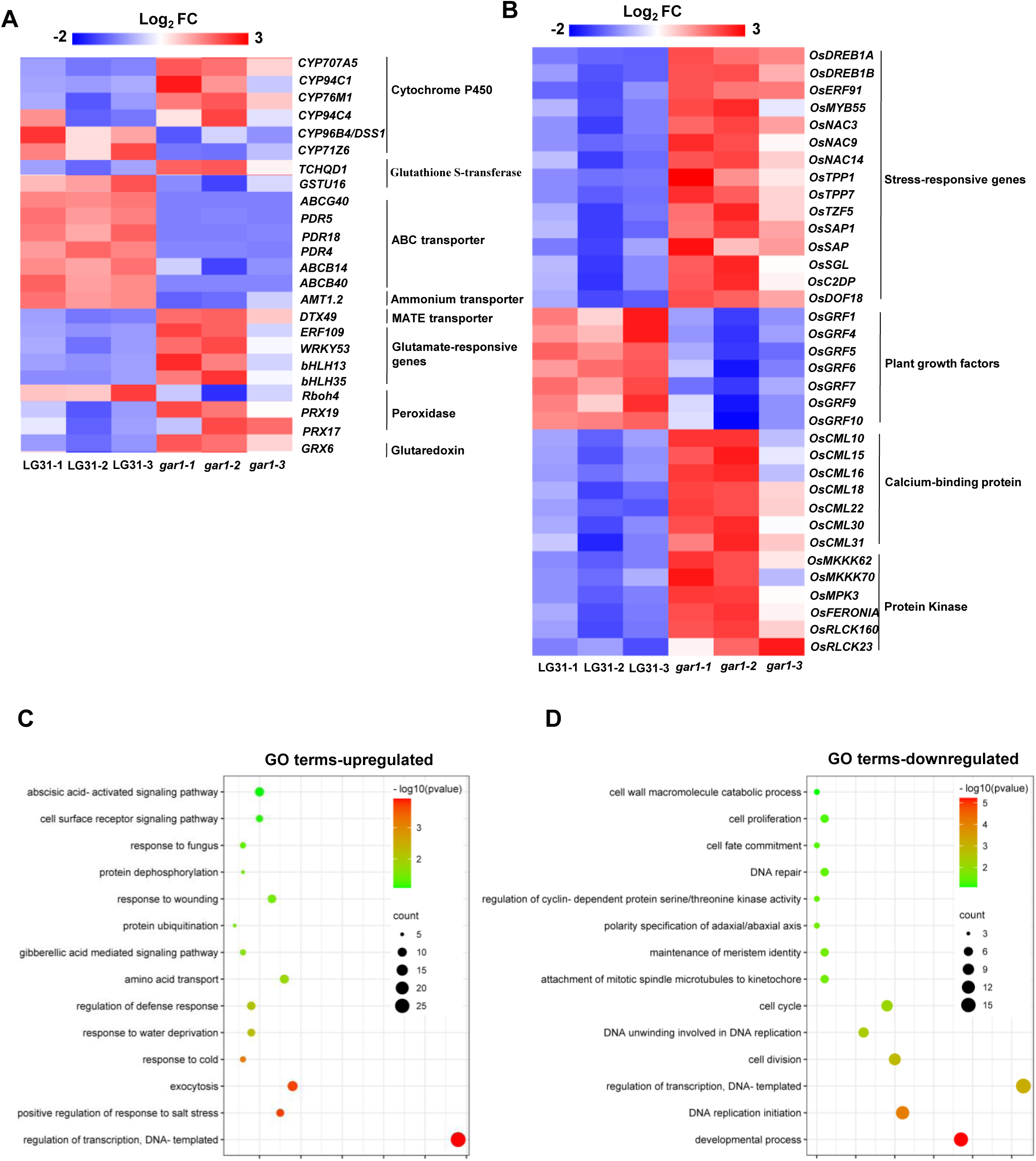
Distribution of differentially expressed genes (DEGs) in LG31 and *gar1-1*. **A-B**. Heatmaps of the enriched possible detoxification-related genes (including cytochrome 450, glutathione S-transferase, ABC transporter, ammonium transporter, MATE transporter, glutamate/glutamine-responsive genes, peroxidase, and glutaredoxin), stress response, plant growth, calcium-binding proteins and protein kinase-related genes are shown between LG31 and *gar1-1*. The fold changes are presented as the log_2_FC ratio. **C-D.** Gene ontology (GO) term enrichment analysis of up– and downregulated genes of LG31 and *gar1-1*. The regulation of transcription, exocytosis, stress response (including salt, cold, water deprivation, defense and fungus) and ABA signaling pathway-related genes were upregulated; in addition, growth and development-related genes (including developmental process, DNA replication initiation, cell cycle and division) were downregulated. The size of the dot plot represents the number of genes enriched in a functional term, and a larger dot area represents a more significant enrichment.

Notably, a large number of regulatory genes related to the stress response were upregulated in *gar1-1*, including *DREB1A*, *DREB1B*, *NAC3*, *NAC9*, *NAC14*, *TPP1*, *SAP*, and *SGL*. However, the expression levels of genes related to plant growth regulation (*GRF1*, *GRF4*, *GRF5*, *GRF6*, *GRF7*, *GRF9*) were repressed in *gar1-1* (Figure 6B). Furthermore, genes encoding calmodulin-like (CML) proteins and protein kinases were also significantly elevated in *gar1-1*. These results indicate that *OsARF18* negatively regulates the expression of the genes associated with the stress response and signal transduction and positively regulates growth regulation genes, which is consistent with the *gar1-1* phenotype of enhanced tolerance to salt and osmotic stress and suggests a role of *OsARF18* in the balance of stress response and growth.

To gain a global view of *OsARF18*-mediated gene expression changes, a gene ontology (GO) analysis was performed for up– and downregulated genes between LG31 and *gar1-1* (Figure 6C, D). In upregulated GO terms, several categories, including response to abscisic acid, fungus, wounding, gibberellic acid, water deprivation, cold and salt stress, were predominantly enriched (Figure 6C), whereas in the downregulated GO terms, DEGs were assigned to developmental process, DNA replication, cell division and proliferation, cell cycle and regulation of transcription (Figure 6D). We also analyzed the DEGs based on the KEGG pathway in depth.

Several categories, including plantLpathogen interaction, zeatin biosynthesis, plant hormone signal transduction, and fatty acid elongation, were enriched in the upregulated group of the LG31 vs. *gar1-1* comparison (Figure. S2B). Meanwhile, the pathways involved in DNA replication, nitrogen metabolism, diterpenoid biosynthesis and ABC transporters were enriched in the downregulated group of the LG31 vs. *gar1-1* comparison (Figure. S2C). These results suggest that loss of *OsARF18* plays an important role in glufosinate resistance and salt and osmotic stress tolerance.

The RNA-seq data were confirmed by RTLqPCR analyses of a number of representative genes, including *Os11g05380* (*P450*), *Os11g05380* (*P450*), *OsDTX49*, *Os09g16458* (*ABC*), *Os09g16449* (*ABC*), *Os02g21350* (*ABC*), *OsDREB1A*, *OsDREB1B*, and *OsTPP7* (Figure S3).

### Improved salt and osmotic stress tolerance of *gar1* mutants

Our transcriptomic analyses indicated that OsARF18 may regulate salt and osmotic stress tolerance. We performed salt and osmotic stress tolerance assays using LG31, *gar1-1*, ZH11 and three *OsARF18* knockout mutants. There were no significant differences among all the lines before salt treatment. After 150 mM salt treatment for 7 days and recovery for 3 days, the *gar1-1* line showed significantly enhanced salt stress tolerance with a survival rate of 86%, which was significantly higher than that of LG31 seedlings (22%). Similar to *gar1-1*, the knockout mutants (*osarf18-1, osarf18-2* and *osarf18-3*) exhibited survival rates of approximately 80%, significantly higher than ZH11 of approximately 20% (Figure 6A-C). These results are supportive of previous reports (Deng et al., 2022).

In our osmotic stress tolerance assay, the *gar1-1* displayed a survival rate of 56% compared with the wild-type LG31 of only 11%. Consistent with this, the survival rates of the knockout mutants (*osarf18-1, osarf18-2* and *osarf18-3*) were 74%, 78% and 86%, respectively, whereas that of the wild-type ZH11 was 33% under the same conditions (Figure 6D-F). These results demonstrated that loss of *OsARF18* also confers enhanced osmotic stress tolerance.

## DISCUSSION

Weed infestation is one of the major factors that greatly affect crop growth and productivity, causing a rice yield reduction of more than 40% (Jin et al., 2022). Herbicides have been widely used in fields for more than half a century as the primary ways of controlling weeds due to their economic and effective measures (Liu et al., 2020). To alleviate the loss caused by weeds, the development of herbicide-resistant crops is an efficient strategy to control weeds in the field and achieve sustainable crop production. In this study, we isolated a glufosinate ammonium resistance mutant, *gar1-1*, and identified the mutation in *OsARF18* (*LOC_Os06g47150*) by whole-genome sequencing, which was confirmed by three additional *OsARF18* knockout mutants generated by the CRISPR/Cas9 system. These results further proved that OsARF18 acts as a negative regulator of glufosinate resistance. In addition, an herbicide dose assay showed that *gar1-1* was at least 3 times more resistant to glufosinate than LG31 (Figure 1B).

To better understand the mechanisms of herbicide resistance in plants for the development of herbicide-resistant crops, scientists have performed much research and revealed several major herbicide resistance mechanisms, including target site resistance (glutamine synthetase) and nontarget site resistance (glufosinate uptake, translocation and metabolism) (Carvalho-Moore et al., 2022; Jalaludin et al., 2017; Meyer et al., 2020; Nazish et al., 2022). *OsGSs*, as known target genes of glufosinate, have been used as major candidate genes for the development of glufosinate herbicide-resistant crops (Avila Garcia et al., 2012; Noguera et al., 2022; Prasertsongskun et al., 2002; Ren et al., 2023; Zhang et al., 2022). Concurrent overexpression of *OsGS1;1* and *OsGS2* improved glufosinate herbicide and other abiotic stress resistance in transgenic rice (James et al., 2018). OsARF18, as a transcription factor, cannot be directly involved in the resistance of the herbicide, which means that it may participate in the resistance of glufosinate by directly or indirectly regulating the expression of target resistance or nontarget resistance genes. Thus, *OsGSs* were first evaluated as candidate target genes for OsARF18, and the results showed that OsARF18 can bind to the TGTCTC/ TGTCNN motif in the promoter region of *OsGSs* and repress their expression, therefore affecting glufosinate resistance (Figure 4 and 5). It is interesting to note that the expression of *OsGS1;1* and *OsGS1;2* was induced by glufosinate treatment only in *gar1* mutants but not in the wild type (Figure 4A, B), indicating that OsARF18 represses this herbicide-induced expression. Therefore, *gar1* mutants gain higher GS activity upon glufosinate herbicide treatment than wild type (Figure 4C).

In addition, cluster analysis results of LG31 and *gar1-1* transcriptome DEGs suggested that *OsARF18* may also be able to mediate glufosinate resistance by regulating other detoxification-related genes, such as CYP450 (*CYP707A5*, *CYP94C1*, *CYP76M1*, *CYP94C4*, *CYP96B4*, and *CYP71Z6*), glutathione S-transferase (*GST3, TCHQD1, GSTU16*), ABC transporters (*ABCG14*, *ABCG40*, *PDR4*, *PDR5*, and *PDR18*), and *DTX49* (Figure 6A). Cytochrome P450s constitute one of the largest superfamilies, are widely present in almost all organisms and have been reported to detoxify multiple herbicides by degradation, such as paraquat, tembotrione, bentazon, and nicosulfuron (Dimaano and Iwakami, 2021; Huang et al., 2022; Powles and Yu, 2010; Siminszky, 2006). ABC transporters or detoxification efflux carrier (DTX)/multidrug and toxic extrusion (MATE) transporters can transport a variety of substances (plant hormones, secondary metabolites and heterologous toxic compounds) and participate in multiple physiological processes (Dahuja et al., 2021; Hwang et al., 2016; Upadhyay et al., 2019). For example, AtPDR11, EcABCC8, and AtDTX6 participate in the uptake, efflux or translocation processes of paraquat or glyphosate (Lv et al., 2021; Pan et al., 2021; Xi et al., 2012; Xia et al., 2021).

Glutathione S-transferases (GST3, TCHQD1, GSTU16) may also be involved in the detoxification of glufosinate (Vaish et al., 2020). Therefore, it is of great significance to further study whether OsARF18 regulates other resistance genes in addition to *GS* genes. Moreover, transcriptome data also revealed that multiple stress response genes and growth-related genes were significantly upregulated and downregulated in *gar1-1*, respectively, suggesting that *OsARF18* may be involved in the regulation of the balance between the abiotic stress response and growth (Figure 6B). Our results also indicate that *OsARF18* negatively regulates osmotic stress tolerance (Figure 7D-F) in addition to salt stress tolerance (Figure 7A-C), which is consistent with a previous report (Deng et al., 2022). Huang et al. showed that the expression levels of *OsARF18* can regulate rice growth and development by affecting auxin signaling, and a high level of OsARF18 can lead to a series of growth and development defects, such as plant dwarfing, fewer tillers, abnormal flower and seed development, and smaller seed size (Huang et al. 2016). However, the molecular mechanisms of how *OsARF18* maintains the balance between the stress response and growth and development remain to be further explored.

**Figure 7.**
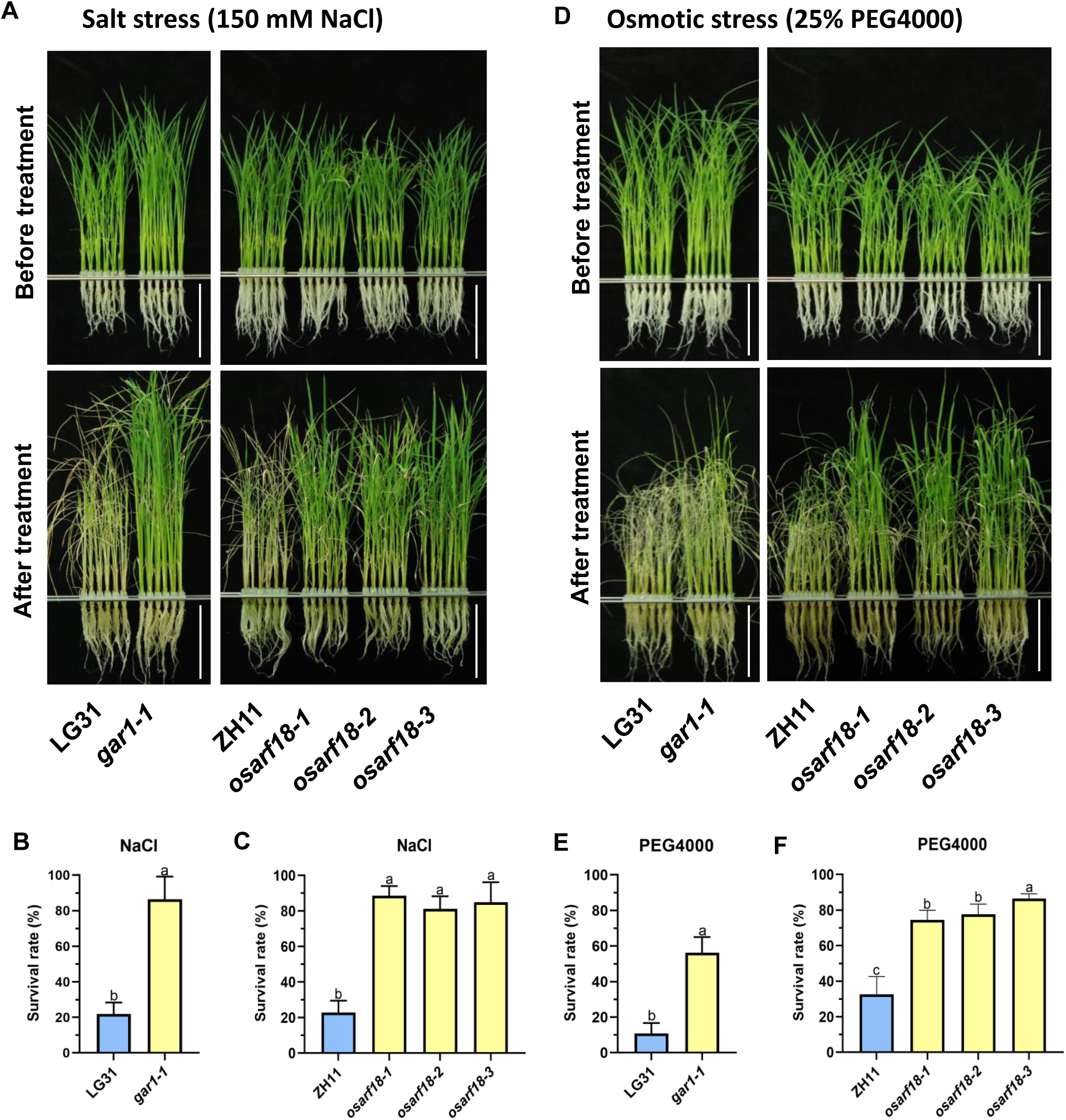
Improved salt and osmotic stress tolerance in *gar1* mutants. **A.** Salt stress tolerance assay. Two-week-old seedlings of LG31, *gar1-1*, ZH11, *osarf18-1*, *osarf18-2* and *osarf18-3* were treated with 150 mM NaCl for 7 days and allowed to recover for 3 days before images were recorded. **B-C.** The survival rates of A. The data in B and C are shown as the means ± SD (n = 3, 48 seedlings per biological replicate). Different lowercase letters indicate significant differences (P < 0.05; one-way ANOVA). Bar = 10 cm. **D.** PEG osmotic stress tolerance assay. Two-week-old seedlings of LG31, *gar1-1*, ZH11, *osarf18-1*, *osarf18-2* and *osarf18-3* were treated with 25% PEG4000 for 2 days and allowed to recover for 5 days before images were recorded. **E-F.** The survival rate of D. The data in E and F are shown as the means ± SD (n = 3, 48 seedlings per biological replicate). Different lowercase letters indicate significant differences (P < 0.05; one-way ANOVA). Bar = 10 cm.

Taken together, our study demonstrated that OsARF18 is a pivotal negative regulator of glufosinate herbicide resistance by regulating GS enzyme activity in rice and is a novel candidate gene for the development of glufosinate ammonium herbicide resistance and stress tolerance crops.

## MATERIALS AND METHODS

### Plant materials and Growth Conditions

Two japonica rice varieties, Longgeng31 (LG31) and Zhonghua11 (ZH11), were used in this study. The *gar1-1* mutant was obtained by screening an EMS-mutagenized rice library in the LG31 background with glufosinate ammonium herbicide at 2 g/L. *OsARF18* knockout mutants (*osarf18-1, osarf18-2* and *osarf18-3*) were generated in the ZH11 background by Hangzhou Biogle Co., Ltd. (Hangzhou, China) (http://www.biogle.cn/) using the CRISPR/Cas9 technique (Lu et al., 2017), and homozygous mutant lines were confirmed by sequencing.

For hydroponic culture, rice seedlings were grown in modified Kimura B nutrient solution in an artificial climate chamber (16-h light/8-h dark, 30°C/28°C cycles.) as described (Wu et al., 2020).

### Genetic screen and identification of glufosinate ammonium resistance mutants

To screen glufosinate ammonium resistance (*gar*) mutants, approximately 1 million EMS-mutagenized M2 seeds in the LG31 background were germinated and grown in soil. Two-week-old M2 seedlings were sprayed with 2 g/L glufosinate herbicide (Ruikai Chemical, Shijiazhuang, China), which was prepared by dissolving 95% commercial glufosinate solids in water. Ten days later, the vast majority of seedlings were withered and yellow, whereas the putative *gar* mutants were still green; then, the putative *gar* mutants were transferred to the field for maturation. M3 seeds were rescreened under the same conditions and back-crossed with wild-type LG31 to generate F1 progeny, and F2 progeny were derived from F1 self-pollination.

Two-week-old F2 seedlings were phenotyped by using the same screening conditions, and 20 F2 segregated resistant progeny seedlings were pooled for whole-genome resequencing by the MutMap method (Abe et al., 2012; Takagi et al., 2015). Genome resequencing was performed commercially by Allwegene Technology Inc. (Nanjing, China).

### Glufosinate spray assay of soil-grown seedlings

To confirm glufosinate tolerance, two-week-old seedlings of LG31, *gar1-1*, ZH11 and *OsARF18* knockout mutants (*osarf18-1, osarf18-2* and *osarf18-3*) in soil were sprayed with 2 g/L glufosinate herbicide as in the *gar* mutant screen. After 10 days, the phenotype was recorded.

### Salt stress and PEG osmotic stress tolerance assays of hydroponic seedlings

To explore salt stress tolerance, two-week-old seedlings were exposed to Kimura B nutrient solution containing 150 mM NaCl for 7 days, and then seedlings were allowed to recover in nutrient solution for 3 days. The survival rates were counted.

For the PEG osmotic stress tolerance test, two-week-old seedlings were treated with 25% PEG4000 for 2 days and allowed to recover for 5 days. The survival rates were counted.

### Subcellular localization of OsARF18-eGFP fusion proteins in transgenic plants and *N. benthamiana*

To investigate the subcellular localization of OsARF18, the CDS of *OsARF18* was fused into the *35Spro:eGFP* empty vector to generate the *35Spro:OsARF18-eGFP* plasmid. GFP fluorescence was observed in the leaf epidermis of *Nicotiana benthamiana* and transgenic seedling roots. GFP imaging was performed by using a Zeiss 980 fluorescence microscope.

### Promoter GUS assay

The 2-kb promoter region of *OsARF18* was amplified from ZH11 and cloned into pCAMBIA2391Z to generate *OsARF18pro:GUS*, and the resulting vector was transformed into ZH11. GUS staining was performed as previously described (Wang et al., 2018).

### Expression pattern of *OsARF18* under abiotic stress

Wild-type seedlings (ZH11) were cultured in modified Kimura B solution for two weeks. Then, the seedlings were treated with 150 mM NaCl, 25% PEG4000, 100 μM ABA, low temperature (4 L), 20 μM H_2_O_2_ and 100 μM CdCl_2_. Rice seedlings were sampled at each time point (0, 1, 3, 6 and 12 h) after treatment, and each group was treated with modified Kimura B nutrient solution as a blank control. For each condition, 6 seedlings were sampled at every time point for RNA extraction, with three biological replications per treatment as previously described (Alfatih et al., 2020; Li et al., 2022c; Liu et al., 2014).

### ChIP**□**qPCR assays

The leaves of two-week-old *35Spro:OsARF18-eGFP* transgenic rice plants grown in soil and GFP-Tag antibodies (Abmart, Shanghai, China) were used for ChIP experiments. Each group contained at least 6 individuals (approximately 1 g) and was treated with the same treatment for wild-type ZH11. The DNA was purified and quantified by qPCR using the primers listed in Table S1.

### Transient trans repression assay

Two-kb fragments of the *OsGS1;1* and *OsGS1;2* promoters were cloned into the pGreen0800 vector to generate reporters. The full-length CDS of *OsARF18* was cloned into the *35Spro:eGFP* empty vector to generate the effector. The reporter and effector were transferred into *Agrobacterium* strain GV3101 (pSoup-p19). Different *Agrobacterium* combinations of equal volume were mixed gently and coinfiltrated into four-week-old tobacco leaves. Plants were grown under 14 h light (28°C)/10Lh dark (25°C) cycles with a relative humidity of 50% for 60 h. Luciferase activity was measured using an assay kit (Lot: 122799-10, PerkinElmer). Images of tobacco leaves were recorded by a CCD system (Feng et al., 2022).

### RNA extraction and real time quantitative PCR (RT**iJ**qPCR) analyses

Total RNA of different samples was extracted by using the TransZol Up Plus RNA Kit (TransGen, Beijing, China), and RNA was reverse transcribed into cDNA with the TransScript Kit (TaKaRa, Dalian, China). RTLqPCR was performed with a StepOne Plus Real Time PCR System by using a TaKaRa SYBR Premix Ex Taq II reagent kit. The specific primers used for RTLqPCR were provided in Table S1.

### RNA sequencing analysis

For RNA-Seq analysis, RNA-Seq was performed by using LG31 (wild type) and *gar1-1* plants. Aerial parts of two-week-old seedlings were harvested for total RNA extraction. Ten seedlings were collected as a sample for each line. and three independent biological replicates were maintained. RNA library construction and sequence analysis were conducted by Beijing Biomarker Technology Company.

### Statistical analysis

Statistically significant differences were computed based on one-way ANOVA.

## Acknowledgments

This work was supported by grants from the Department of Science and Technology of Anhui Province (grant no. 202003a06020027) and the China Postdoctoral Science Foundation (2022M723053). We thank Prof. Chengcai Chu (South China Agricultural University, Guangzhou, China) for providing the pCAMBIA2391z vector and Dr. Yangwen Qian (Biogle Genome Editing Center, Changzhou, China) for providing the *35Spro:eGFP* vector.

## Author contributions

CBX and JQX designed the experiments. DYH and JQX performed most experiments, and JQX and PXZ analyzed the data. JQX, QYL, JW, ZSZ, JZ and ZYZ participated in genetic screening of the *gar* mutants or assisted in part of the molecular experiments. DYH and JQX wrote the manuscript. CBX revised the manuscript and supervised the project.

