## Supplemental info for "Loss of *OsARF18* confers glufosinate ammonium herbicide resistance in rice"

### Supplemental information for "Loss of *OsARF18* enhances glufosinate ammonium herbicide resistance in rice" by He et al.

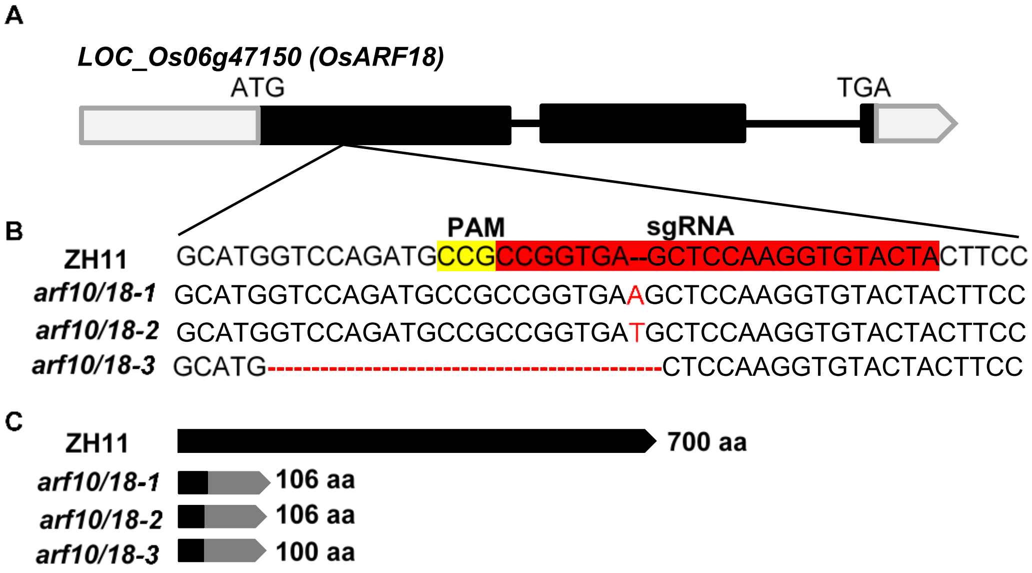

**Figure S1. Identification of CRISPR/Cas9-edited *OsARF18* knockout mutants**

**A.** Schematic representation of the *OsARF18* gene. Black boxes represent exons, gray boxes represent UTRs, and black lines represent introns.

**B.** Sequence around the CRISPR/Cas9 target site (sgRNA) of wild type (ZH11) and the three knockout mutants (*osarf18-1*, *osarf18-2* and *osarf18-3*). PAM, protospacer adjacent motifs. sgRNA, single guide RNA.

**C.** Truncated OsARF18 protein in *osarf18-1*, *osarf18-2* and *osarf18-3*. The black box indicates the amino acid sequence translated correctly, and the gray box indicates the amino acid sequence translated after the frameshift.

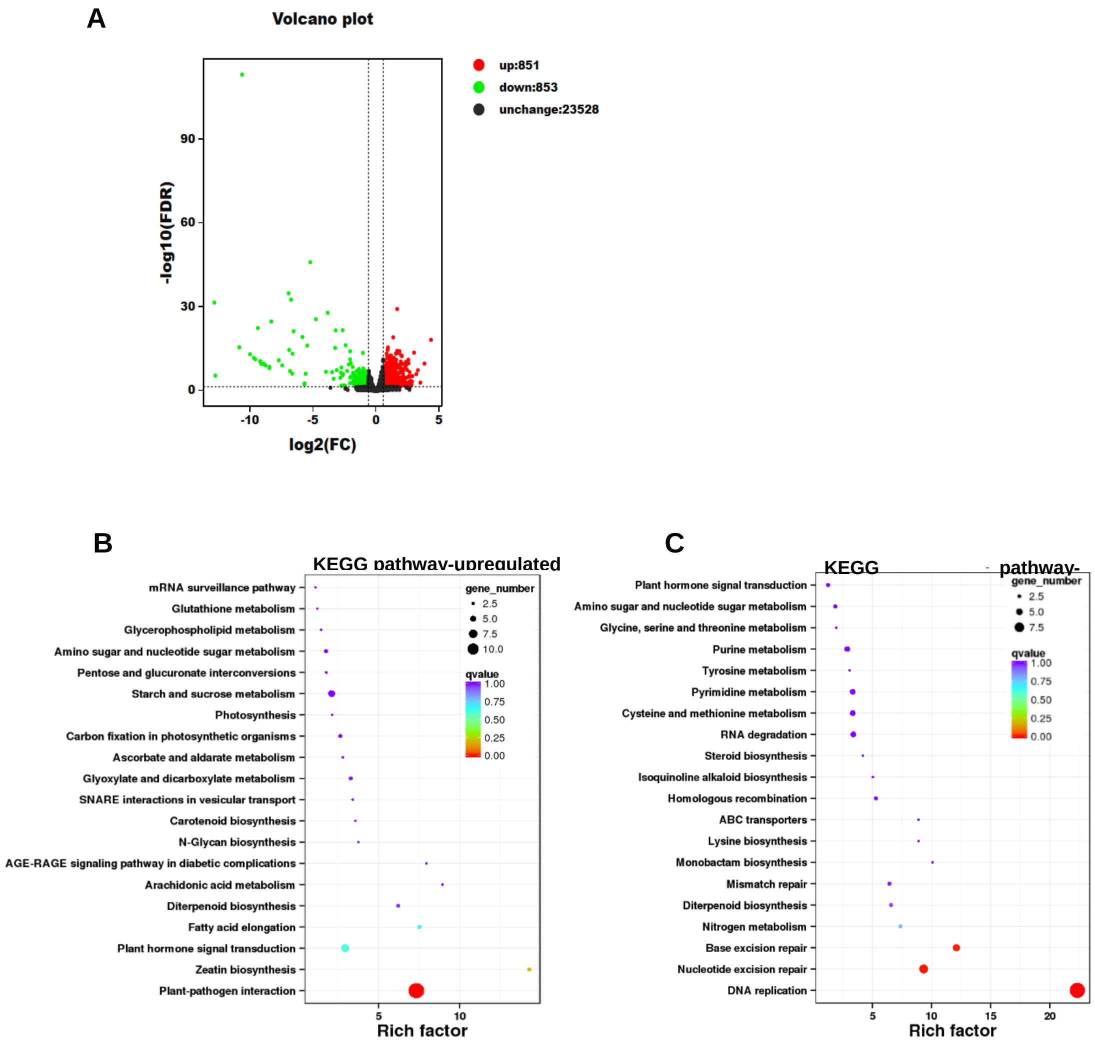

**Figure S2.** Differentially expressed genes (DEGs) between LG31 and *gar1-1* and KEGG pathway analysis.

**A.** Volcano plot of DEGs. The X-axis represents the value of the log2 (fold change). The Y-axis represents the negative log10 value of the FDR. Red dots indicate upregulated genes, green dots indicate downregulated genes, and black dots indicate unchanged genes.

**B-C.** KEGG pathway analyses of up- and downregulated genes of LG31 and *gar1-1*.

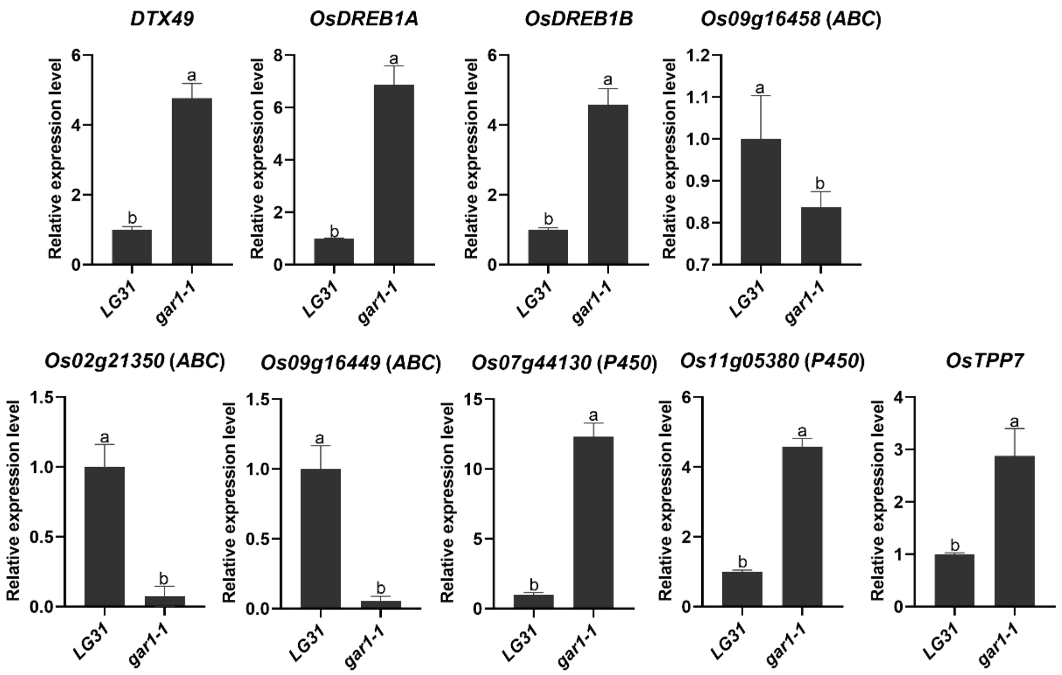

**Figure S3.** Validation of representative up- and downregulated DEGs in LG31 and *gar1-1* by RT–qPCR

**Table S1.** Primers used in this study.

| Primer | Forward sequence (5'-3') | Forward sequence (5'-3') |
|---|---|---|
| OsACTIN-Q-PCR | CACAGGTATTGTGTTGGACTCTG | AGTAACCACGCTCCGTCAGG |
| OsGAR1-Q-PCR | CTTAACAAACTGCAGTCGAGTC | CTGATTTCTTGTTGTTCTGGGG |
| OsGS1;1-Q-PCR | GAAGGGAAACAACATCCTTGTC | CGATACCACAGTAGTAAGGACC |
| OsGS1;2-Q-PCR | GAAGTGATCCTCTACCCTCAAG | CCTCTTGTTAGTGGGGATTGG |
| OsGS1;3-Q-PCR | AGCCGATAAATCGTACGGGC | AATCTGGAACTCCCACTGCC |
| OsGS2-Q | CGTCGGGGTTCAGGGTGATG | GGAGCTTGCCCTGTGCTTGA |
| OsDTX49-Q-PCR | GGCGGAGGTCGTCATCTTGG | AGCAGGTGAACATGGACGGC |
| OsDREB1A-Q | GACGTCCTGAGTGACATGGG | AGTAGCTCCAGAGTGGGACG |
| OsDREB1B-Q | ACGATGGCGACGAAGAAGAAGACA | TGCGCCAAGCTCGCGTAGTA |
| OsDREB1C-Q | CGCCCGCCATGATGATGCAGTA | ATCGTCGTCGCCGTCCATCT |

| | | |
|---|---|---|
| OsDREB1E-Q | CTTCCCTTGCTACCCGATGG | TTGACCTCGCAGTCGTAGTC |
| OsDREB1G-Q-P1 | CGCCACTAATTCGAACGCC | TGTCGAGATAGCCCTGCATC |
| P450 (Os07g44130)-Q | ATGGTGTGCAATGACGAGGT | TTGCGATCGGGATCGTTAGG |
| P450 (Os11g05380)-Q | CTTCTTCGCGTTCTCGGTGT | GAGGTTGGTGAACTCCCTCG |
| OsTPP7-Q | TGCGTCGACGAGAAGAGTTG | CCCTTGTCCCACTTGATGGT |
| pHIS-GS1;2-ARF18-CIS1 | AATTCCTGAAAAAAAAATGTCTCTGAATGATACA | CGCGTGTATCATTCAGAGACATTTTTTTTCAGG |
| pHIS-GS1;2-ARF18-CIS2 | AATTCGCCACGGCCGTTGTCGGATTTAACATTTA | CGCGTAAATGTTAAATCCGACAACGGCCGTGGCG |
| pHIS-GS1;2-ARF18-CIS | AATTCCAACAATGATTGGAGACATGTAACCAGGA | CGCGTCCTGGTTACATGTCTCCAATCATTGTTGG |
| pGreen0800 | TAAGTTGGGTAACGCCAGGG | CTCTCCAGCGGTTCCATCTT |
| CHIP-GS1;1-ARF-CIS | TGTTACAGCTGTGCCGCCT | AACCAAGAGCCAAGAAGAGATCC |
| CHIP-GS1;1-ARF-CIS1 | TGCTTTCCTCCCACGTAGAAACC | GTATGTGATGAATAGAATAGCACTCTCTCC |
| CHIP-GS1;1 | ACGACAAAGAACAGGGAGTGG | GGGAAAAATTATTTATCCCGAGCGGC |
| CHIP-GS1;2-ARF-CIS1 | ACACTTAAGACGTTGGGATTAGCAA | TTCTTCTCTTCCCGGCCTGTTG |
| CHIP-GS1;2-ARF-CIS2 | GCTTGTTTATGCTTAGACAAACTCTCAA | GTTCGTCACATTTCTGCTCTGTATTT |
| CHIP-GS1;2-ARF-CIS | ACCAGAGGTAATCCATTCCCTTGA | AAGTTAGCTACGGTGTGCTTGCAA |
| CHIP-GS1;2 | GGAGCATCACGGCCAAATTCT | GCTTGCACATTCACGTTTATTAACTT |

| | | |
|---|---|---|
| ABC (Os09g16449) | GTTCGGAACCATGTTCTGGG | AAGATCGTGCGCCTCAACAA |
| ABC (Os09g16458) | GCCCTCGTGATCATGTGCTT | CCATGGTAAAACTCACTTCCTGC |
| ABC (Os02g21350) | GGTGATGACGACCACAAGGA | TTCTGGCTCCGCCACTATAC |
| pHis-OsGS1;2-arf1 | CTGATTGGAGACATGTAACA | CGCGTGTTACATGTCTCCAATCAGAGCT |
| pHis-OsGS1;2-arf2 | CAAAAAATGTCTCTGAATGA | CGCGTCATTCAGAGACATTTTTTGAGCT |
| pHis-OsGS1;1-arf1 | CCAAAGATGTCTCATAGGCA | CGCGTGCCTATGAGACATCTTTG |
